# An easy-to-use microfluidic mechano-chemostat for tissues and organisms reveals that confined growth is accompanied with increased macromolecular crowding

**DOI:** 10.1101/2023.03.29.534752

**Authors:** Zacchari Ben Meriem, Tiphaine Mateo, Julien Faccini, Céline Denais, Romane Dusfour-Castan, Catherine Guynet, Tatiana Merle, Magali Suzanne, Mickaël Di-Luoffo, Julie Guillermet-Guibert, Baptiste Alric, Sylvain Landiech, Laurent Malaquin, Fabien Mesnilgrente, Adrian Laborde, Laurent Mazenq, Rémi Courson, Morgan Delarue

## Abstract

Conventional culture conditions are oftentimes insufficient to study tissues, organisms, or 3D multicellular assemblies. They lack both dynamic chemical and mechanical control over the microenvironment. While specific microfluidic devices have been developed to address chemical control, they are often hard to use and do not allow the control of compressive forces. Here, we present a set of microfluidic devices which all rely on the use of sliding elements consisting of microfabricated rods that can be inserted inside a microfluidic device. Sliding elements enable the creation of reconfigurable sealed culture chambers for the study of whole organisms or model micro-tissues. By confining the micro-tissues, we studied the biophysical impact of growth-induced pressure and showed that this mechanical stress is associated with an increase in macromolecular crowding, shedding light on this understudied type of mechanical stress. Our mechano-chemostat is an easy-to-use microfluidic device that allows the long-term culture of biological samples and can be used to study both the impact of specific conditions as well as the consequences of mechanical compression.

## Introduction

Cells in tissues and organisms, or during development, are constantly subjected to dynamic chemical and mechanical cues. Imposing dynamic chemical conditions on 3D cellular assemblies is a technical challenge which requires the use of complex microfluidic devices^1–4^. However, and despite the large parallelization enabled by some of these devices, they do not necessarily allow easy dynamic control, and very few enable the establishment of chemical spatial gradients^5,6^ which are essential to study 3D chemotaxis or drug screening. Mechanically, and apart from devices allowing a control of shear or tensile stresses^7,8^, the appropriate 3D mechanical conditions to study the effect of spatial confinement and compressive stresses are lacking.

Compressive stresses can either be dynamic, such as peristalsis during digestion or the compression of articular cartilage during motion^9^, or self-inflicted in the case of spatially constrained growth^10^. Indeed, compressive stress naturally arises when cells proliferate in a confined space, like solid tumors growing within an organ^11^. Compressive stresses can be deleterious for tumor treatment since they can clamp blood vessels^12^, modulate cell proliferation^13–15^, and even participate in a mechanical form of drug resistance^15^. In contrast with tensile and shear stresses^16–21^, very little is known about the sensing of mechanical pressure.

Growth-induced pressure is notoriously hard to study, all the current methods to impose spatial confinement rely on spheroid embedded in a hydrogel^13–15^. Such methods display natural limitations in terms of the type and size of the studied sample alongside its retrieval for further biological characterization and the dynamic control of the culture conditions.

In general, the culture of organisms inside microfluidic devices remains difficult to do, even though microfluidic systems can offer much tighter control than classical culture. In this paper, we present a set of microfluidic devices that take advantage of an innovative technology we called sliding elements. Sliding elements are microfabricated rods that can be inserted inside a microfluidic device. Using this technology, we created reconfigurable easy-to-use culture chambers which could be loaded with biological objects such as spheroids or living organisms. These devices permit great chemical control, real-time imaging and the possibility to recover the sample. Moreover, we enforced a tight mechanical control over the sample, with the possibility to either dynamically compress it, or to spatially confine it in order to study the impact of growth-induced pressure. Hence, novel pressure sensors have been developed to measure mechanical pressure. We demonstrated the great versatility of our mechano-chemostats through the long-time culture of spheroids of different cell types, drosophila legs and nematodes. We showed in particular that growth-induced pressure was associated with increased macromolecular crowding, thus shedding light on a novel biophysical regulation of confined growth in mammalian cells.

## Results

### Sliding elements to create a microfluidic chemostat for biological samples

The realization of a microfluidic chemostat resides in our ability to load a sample at a given position and define the chemical environment around it (Fig. 1a). Valves could be used to trap a sample, but the feeding remains difficult. Solutions relying on one-way valves have been developed for microbes^10,22^, but are not directly amenable to larger and deformable samples. To overcome this difficulty, we underwent a key technological development: sliding elements, tiny 3D-structured rods which can be inserted inside a microfluidic system to bring specific functions of interest^23^. By coupling standard photolithography and the use of dry film photoresists, we created well-defined and transparent sliding elements with cylindrical holes or slits depending on the direction of fabrication (Fig. 1b). They were centimetric in length and squared in the other dimensions with a cross size of 500μm, making them easy to manipulate and slide into a designated channel (Fig. 1c). We created them by the hundreds in one batch (Fig. 1c, inset).

**Figure 1:**
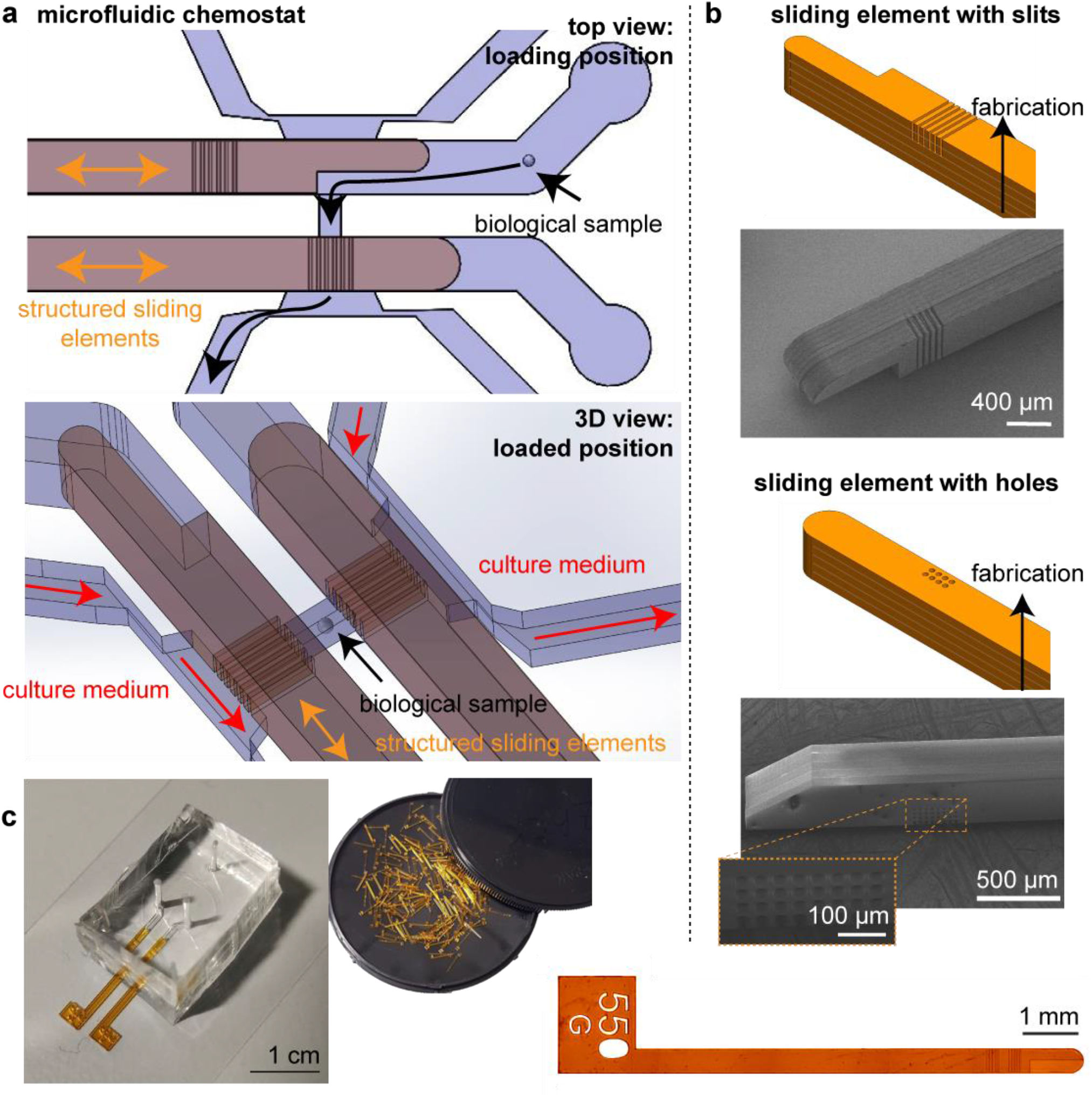
Design of the microfluidic chemostat. **a**. The microfluidic chemostat is composed of a culture chamber which is closed on both sides by structured sliding elements. These elements enable to load the chamber, and feed it thanks to channels on both sides. **b**. Standard photolithography is used on dry films to structure in 3D the element. Depending on the direction of construction, we can either construct slits or holes. Scanning electron images of the sliding elements are presented. **c**. Picture of the microfluidic device with the sliding elements inserted. The sliding elements are centimetric in length and structured at the tens of micrometer resolution. They are fabricated by the hundreds, and can be inserted in a PDMS chip.

Culture chambers were molded in polydimethylsiloxane from molds created using multi-level photolithography, the first one defining the height of the culture chamber, while the second one delineated the channel into which the sliding element would be inserted (Fig. 1c). The height of this channel had to be optimized to ensure tight sealing and avoid medium leakage from one compartment to the next. We find that a PDMS channel of the same size ± 10μm of the size of the sliding worked without leakage for fluid pressure below 200 kPa, which was above the typical maximum 50 kPa pressure needed in our experiments to culture cells. This tight sealing was essential to enable a perfect control over the chemical environment. Notably we showed that we could instantaneously change the chemical conditions in the chamber (Fig. S1). We could have fresh medium with constant chemical conditions circulating or allow a fix volume of medium to cyclically re-circulate in the chamber to either decrease waste or perform specific enrichment experiments.

### Steady culture of multicellular spheroids, moving organisms and imaginal disks

The chemostat could be easily loaded with various biological tissues or organisms. Sliding one element down opens one side of the chamber, so that by adjusting the inlet flow, we could control the position of a multicellular spheroid inside the chamber, pushing it to the end, or retrieving it. We showed that spheroids can be cultured in the device for days (Fig. 2a and supplementary video S1), with no significant differences in growth measured inside the device in comparison with classical culture in well plates (Fig. 2b). Of note, we could parallelize the chambers, different spheroids could be loaded in different chambers (Fig. 2c), or two different samples in the same chamber (Fig. 2di-ii). While the former device showed increase in throughput of the device or parallelized experiments, the latter could be interesting to study interactions (mechanical and chemical) between different samples.

**Figure 2:**
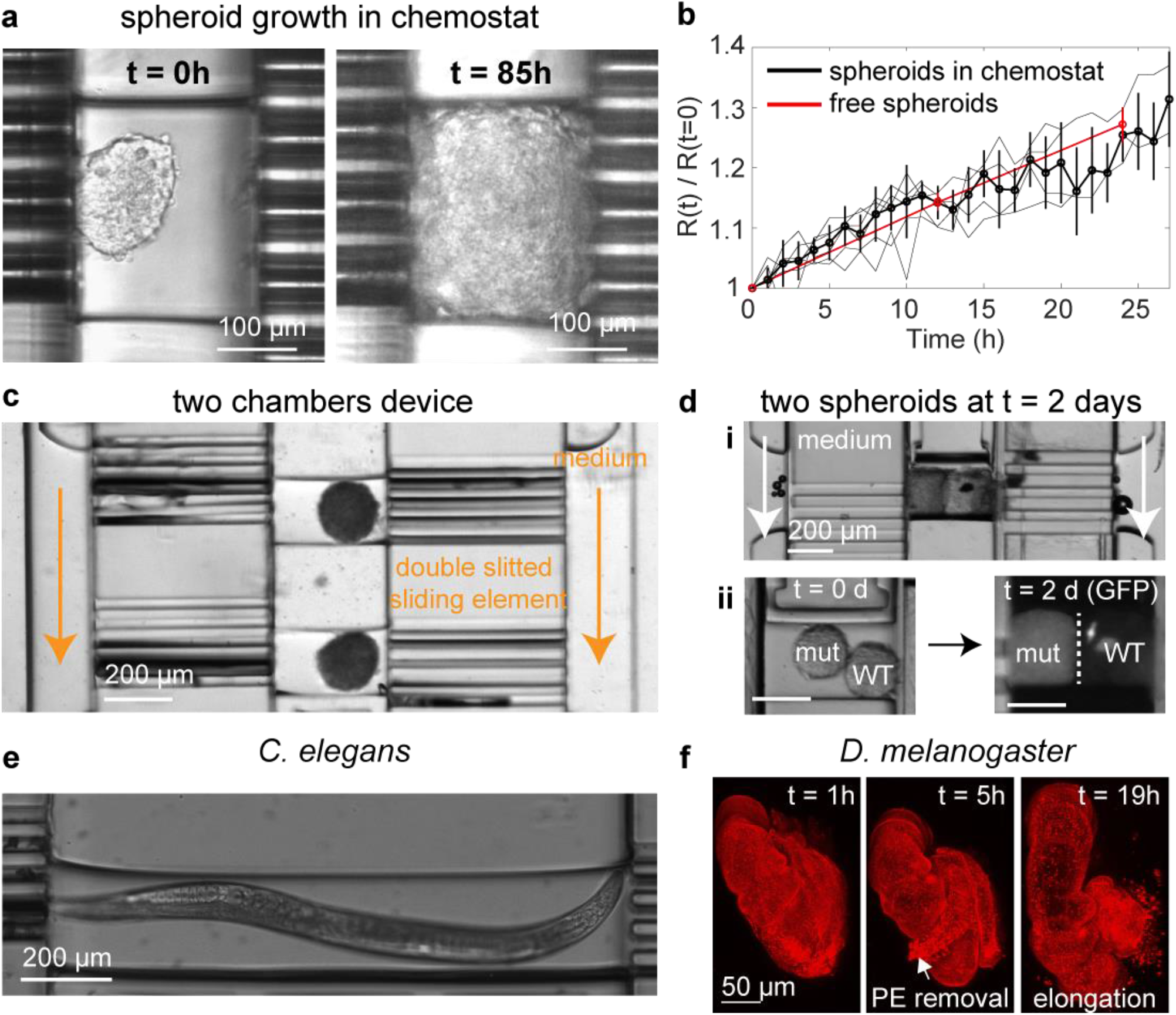
Culture of biological samples in the microfluidic chemostat. **a**. Multicellular spheroids can be loaded in the chemostat. They can grow until they fill the chamber. **b**. Growth curves of spheroids in the chemostat (6 independent replicates in light black) and in classical round bottom well plates (mean ± SEM). Thick lines represent median ± standard deviation. **c**. We designed devices with two parallel chambers where different samples can be loaded and cultured. **d**. Two different spheroids can be loaded and cultured in the same chamber (i). They grow until the chamber is filled (ii). **e**. Moving samples such as the nematode C. elegans can be cultured in the device. **f**. Imaginal disks such as a drosophila leg can be injected, and display normal development in the microfluidic chemostat, as seen by the timing of PE removal and leg elongation.

Our ability to load a biological sample in a closed environment was useful for moving organisms which could be hard to image. As an example, we showed that we could harmlessly load the nematode *C. elegans* and culture it for hours (Fig. 2e and supplementary video S2). The worm remained trapped in the culture chamber, permitting its imaging for hours under fixed chemical conditions.

Biological samples can be cultured for hours / days in the microfluidic chemostat, in a controlled chemical environment. We validated the loading and culture of imaginal disks, such as the *Drosophila melanogaster* leg (Fig. 2f, supplementary video S3). The easy manipulation and culture in the chamber allowed to monitor its development which was similar in the chemostat compared to classical culture conditions^24^. The steady chemical environment, produced using syringe pumps, allowed long culture times, typically hard to reach with classic culture conditions where culture medium volume is fixed^25^.

Importantly, the chamber can be re-opened, and the sample retrieved for further biological analysis. This essential point was often a bottleneck in microfluidics, which the use of sliding elements easily overcame.

### Confined proliferation and growth-induced pressure

While these chemostats revealed to be adequate for the study of specific questions requiring a culture chamber with controlled chemical conditions, they were limiting for the study of the impact of spatial confinement and compressive stresses. Indeed, they did not fully confine a multicellular spheroid, as we observed that, after reaching 3D confluence, the spheroid leaked into the feeding channels, and grew into them in hours. Cells are indeed able to migrate and deform through constrictions as small as 5μm^26^, which was a resolution not reachable during sliding element fabrication. To overcome this issue, we designed a novel three-layer system with a culture chamber connected on its side to much smaller channels (2μm x 2μm in cross section) which fully blocked the spheroid (Fig. 3a). We adapted the design of the sliding element to load and close these chambers (Fig. 3b and supplementary video S4), and observed that spheroids grew fully confined in this geometry (supplementary video S5), without invading the side channels. Normal growth of the spheroid was measured before being spatially confined (Fig. 3c), suggesting optimal feeding.

**Figure 3:**
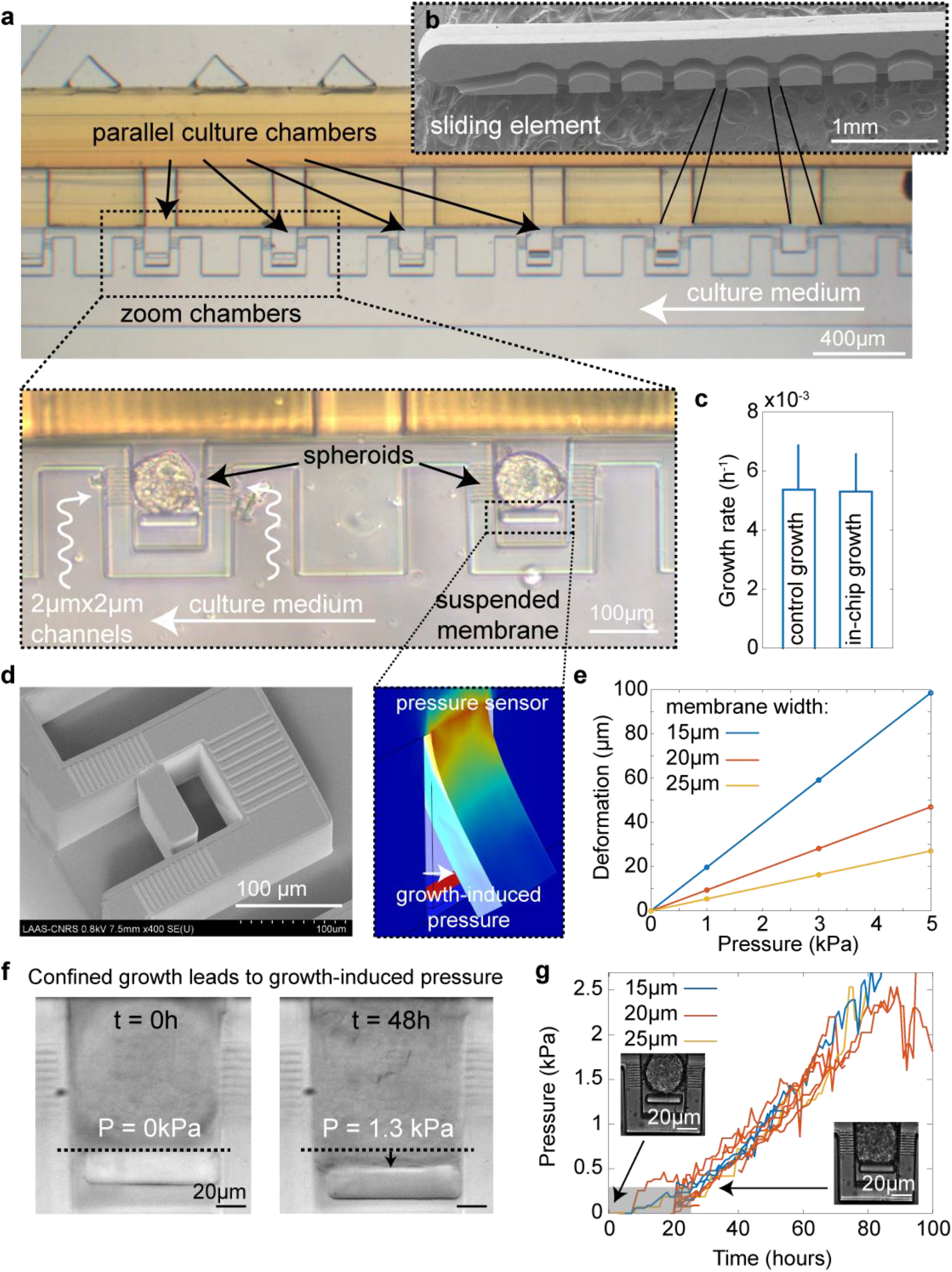
Confined growth of multicellular spheroids and pressure sensor. **a**. The design can be parallelized and built on three levels to create multiple closed culture chambers. **b**. The sliding element is structured in such a way to allow loading and closing of the chambers. **c**. Growth rate of multicellular spheroids before confinement is similar to that of free spheroids (median ± standard deviation, N = 4 independent experiments). **d**. Scanning electron microscopy image of the chamber containing the suspended membrane. Image of a finite element simulation showing its deformation when a fixed pressure is applied. **e**. Deformation of the membrane with applied pressure as a function of membrane width. **f**. Confined growth leads to growth-induced pressure measured by the deformation of the suspended membrane. **g**. Pressure is independent on the width of suspended membrane. After a slow increase which corresponds to a change of spherical shape to a cube, pressure increases roughly linearly for hours. The gray area corresponds to the time points for which pressure is underestimated owing to the aggregate not fully contacting the surface. 10 spheroids over 4 independent experiments.

Confined growth eventually leads to the buildup of growth-induced pressure^27^. Evaluating growth-induced pressure often relies on the measurement of the surrounding deformation^13,15,28^, or the deformation of exogenous sensors such as hydrogel beads^29,30^. Alternatively, micropillars have been widely used to measure kPa stresses exerted by moving cells^31^ or growing spheroids^32,33^, due to their high deformation when sufficiently thin. We adapted this technology to design a thin suspended membrane to measure growth-induced pressure (Fig. 3d). We performed finite element simulations to tune its dimensions to be sensitive to the kPa range^15^ (Fig. 3e). We observed that at similar dimensions, a fully attached membrane was much less deformable than one attached only at the top (Fig. S2). In order to calibrate the mechanical properties of the PDMS, crucial parameter to perfectly infer the pressure exerted onto the membrane from its deformation, we designed a fully attached membrane and measured its deformation with a fixed pressure. This deformation can be compared to finite element simulations to properly calibrate the mechanical properties (Fig. S3). Moreover, we could also use this membrane to instantaneously compress a trapped multicellular spheroid or a collagen gel (Fig. S4).

We observed that confined proliferation of a spheroid led to the progressive build-up of growth-induced pressure over the kPa range in several days (Fig. 3f and supplementary video S5). The dynamics did not depend on the width of the suspended membrane (Fig. 3g), and was very comparable to what would be expected for these cells using a standard hydrogel embedding (Fig. S5). This indicated that cells were similarly fed in both conditions and that growth-induced pressure development did not depend on the type of spatial confinement. Note that we needed to apply a correction factor when the spheroid did not fully contact the membrane (Fig. S6). Because this factor could not be easily determined with our imaging conditions, for pressures below 250 Pa, the pressure was underestimated – these points were grayed on the figure. Interestingly, we observed that during the first 24h, the spheroid deformed into a cuboid, while developing a growth-induced pressure of ∼ 300 Pa. We showed (see Methods) that this information can be used to quantify the surface tension of a spheroid, which in this case is in the range of 1.5 mN/m, consistent with measurements in other cell types done with classical micropipette aspiration^34^.

### Growth-induced pressure increased intracellular crowding and decreased proliferation

We sought to investigate the cellular response to growth-induced pressure. We measured a cellular densification within the compressed tissue, suggesting that single cells were more compressed under confined growth (Fig. S7). Taking advantage on the fact that microfluidics allows high-resolution imaging, we used the FUCCI cell-cycle marker (Fig. 4a) and measured a progressive accumulation of G1 cells as growth-induced pressure increased (Fig. 4b). This result was consistent with former findings showing an association between growth-induced pressure and physiological changes, and notably a decrease in cell proliferation^13,15,28,35,36^.

**Figure 4:**
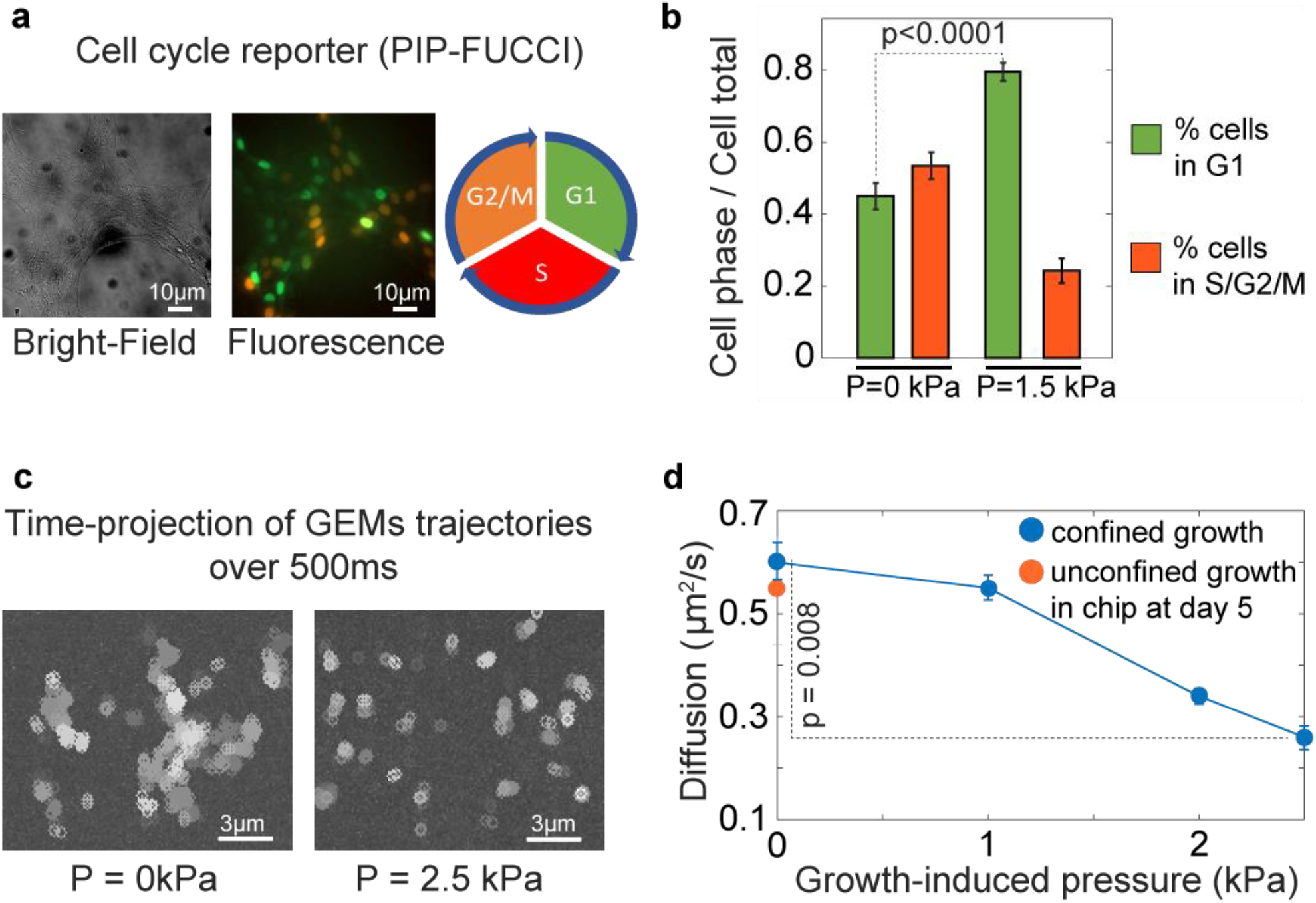
Confined growth lead to growth-induced pressure which impacts cell proliferation and intracellular crowding. **a**. FUCCI cell cycle reporter to fluorescently label cell cycle phases. **b**. Cells accumulate in G1 as growth-induced pressure builds up. 6 spheroids over 4 independent experiments. **c**. Time projection of GEMs nanoparticles trajectories show that particles are less diffusive under growth-induced pressure. **d**. Diffusion progressively decreases as growth-induced pressure increases. N ≥ 10 cells for each point coming from 6 spheroids over 3 independent experiments. For all points, we computed mean ± standard error of the mean.

An elusive question in mechano-biology relates to how growth-induced pressure is integrated and, especially which cellular biophysical properties are modified. It has recently been shown in the budding yeast *Saccharomyces cerevisiae* that growth-induced pressure is accompanied with an increase in intracellular crowding^27^, which relates to the high packing fraction of macromolecules in cells^37^. Genetically-encoded multimeric nanoparticles (GEMs) can be imaged at the single cell level in order to infer intracellular crowding through single particle tracking^38^ (Fig. 4c). Using GEMs, we sought to investigate how intracellular crowding was modified in mammalian cells during the buildup of growth-induced pressure. We found that the mean diffusion coefficient was decreasing with increased growth-induced pressure (Fig. 4d), suggesting that, similarly to *S. cerevisiae*, intracellular crowding increased during confined proliferation and with the buildup of growth-induced pressure.

## Conclusions

We reported here a set of microfluidic chemostats allowing the culture of biological tissues, whole organisms or imaginal disks. Their operation relied on a key and novel technological development, sliding elements, which could be inserted inside a PDMS device to create reconfigurable culture chambers. Sliding elements could be produced by the hundreds, and allowed exquisite resolution thanks to the power of photolithography. In particular, they could be structured by channels or holes, which allowed us to close a culture chamber while retaining the ability to feed the sample loaded in this chamber, something that a classical valve could not do.

Using these sliding elements, we were able to culture different multicellular organisms, from a growing multicellular spheroid to a developing imaginal disk or a moving nematode. Their growth and development were similar to those outside the culture chamber, demonstrating their correct feeding. Additionally, the ability to culture a moving organism in a well-defined position or an imaginal disk for long times, with a dynamically-controlled chemical environment, opens novel avenues in the field of developmental biology. In particular, long term and controlled feeding of imaginal disks allows for the examination of later developmental stages which are, to our knowledge, extremely hard to attain in steady culture conditions.

We showed that we could close the chambers to a point of complete spatial confinement. Vertical membranes can be used to compress the sample. Confined growth of multicellular spheroids led to the buildup of growth-induced pressure, which has a number of physiological consequences. We developed a novel mechanical sensor to measure mechanical pressure, and demonstrated that spheroids in our device could develop growth-induced pressure. In particular, their transition from a spheroid to a cuboid shape allows the estimation of the tissue surface tension independently of other viscoelastic and poromechanics parameters. How growth-induced pressure is integrated and impacts cells is mostly unknown, in contrast to other types of mechanical stresses, such as tensile^16^ or shear^21^. We showed that while cell proliferation was decreased, intracellular crowding increased concomitantly with growth-induced pressure in mammalian cells, yielding a novel biological insight on the mechanisms that can be associated with the integration of growth-induced pressure.

In conclusion, we developed a set of single cast microfluidic devices for the long-term culture of biological samples. These devices are easy-to-use, parallelable to increase throughput, and can be used to study both the impact of specific conditions and the consequences of mechanical compression as well as mechanically characterizing a multicellular spheroid. Compressive stress is still poorly understood owing to the lack of tools available to researchers. Our device offers an elegant solution to its study.

## Author contribution

ZBM, TM, BA and MD designed the culture chambers. ZBM, TM, BA, LM, RC and MD designed the sliding elements. FM, AL, LM and RC helped with microfabrication. TM, CD, MDL, JGG and MD developed cell lines and performed spheroids experiments. ZBM, RDC, CG and MD performed the *C. elegans* experiments. ZBM, TM and MS performed the *D. melanogaster* experiments. SL and MD developed the mathematical analysis. ZBM, TM and MD wrote the manuscript. All authors brought corrections to the manuscript.

## Acknowledgement

The authors would like to thank B. Venzac for critical reading of the manuscript. This work was partly supported by the French Renatech network. MD would like to thank Inserm Plan Cancer (Press-Diag-Therapy and MechaEvo grants), INCa PLBIO and Cancéropôle Grand Sud-Ouest. This work is partly funded by the European Union (ERC, UnderPressure, grant agreement number 101039998). Views and opinions expressed are however those of the author(s) only and do not necessarily reflect those of the European Union or the European Research Council. Neither the European Union nor the granting authority can be held responsible for them.

## Material and Methods

### Device microfabrication

The chemostat is made from a two-layer silicon mold. The high-throughput tumor-on-chip is made from a three-layer silicon mold. For the high-throughput device, we have an initial layer allowing to create the culture channels. This layer is not present in the chemostat where feeding is ensured through the sliding element. All layers are created using dry film technology.

In order to generate channels alimentation which are characterized by a very tiny section of 2×2μm, an initial layer made of a mix of two SU8 photoresist (SU8-6000.5 and SU8 60005, ratio 1:1) is spin-coated (speed: 2500rpm, acceleration: 3000rpm/s, time: 30s) with the spin coater Suss Microtec, on a silicon wafer substrate and cured (2min at 100°c). The photoresist is exposed with the MA6 Gen4 machine (I-line 37% at 300mJ/cm^2^) with the first mask design and cured (100°C during 2min) by following standard photolithography processes. To create the second layer defining the culture chamber, a 100μm dry film is laminated above the mold (pressure: 2.5bars, speed: 0.5m/min, temperature: 100°c for all lamination), and is exposed using a second mask (I-line 37% at 240mJ/cm^2^) and cured (100°c during 6min). The last layer is created from a stack lamination of four 100μm dry-film sheets in order to create the 500μm channel used to insert the sliding element. Then, exposure is performed (I-line 37% at 2000mJ/cm^2^) and the mold is cured (PEB of 100°c during 20min). During exposure steps, particular caution is necessary to align each level with the previous one.

A chemical development in SU8-developper bath is done at the end of the process in order to reveal the channels. Afterwards, a hard-bake is performed in order to reinforce the mechanical resistance of the mold through time. A perfluorodecyltrichlorosilane (FDTS) self-assembled monolayer is grafted onto the surface to prevent polydimethylsiloxane (PDMS) adhesion.

PDMS is cast onto the mold and cured at 65°c overnight. The chip is initially sealed with a thin 50 μm PDMS layer by plasma activating the two surfaces with oxygen plasma (0.2mBar, 0.7sccm, 25s) with the Diener Electronics machine in order to have the same material onto the culture chambers walls. Finally, the whole chip is sealed on a glass slide using same parameters for plasma O2 activation.

Once made, the mold surface is controlled by Scanning Electron Microscopy (MEB Hitachi S-4800). Tension and current are respectively set at 0.6kV and 8μA. To correct astigmatism, magnification is set at x3000. Image definition is about 1200×900px.

### Sliding element fabrication

The sliding element is made of two different levels (300μm and 200μm), using dry film technology, which allows additive fabrication. Each level required stack lamination of 100μm dry film sheet and are laminated using same parameters as the mold fabrication. Starting from a silicon wafer substrate, three dry film of 100μm are successively laminated on it. This one is exposed with a first mask (i-line HR 66mW at 1400mJ/cm^2^) and cured (6min at 100°c) by following standard photolithography processes. The second level is made from two successive lamination of 100μm dry film sheets. Insolation is done using the second mask (I-line HR 66mW, 900mJ/cm^2^) and the mold is finally cured (100°c for 3min). While performing the development bath overnight in SU8 developer, all the sliding elements progressively detach from the wafer substrate, as no adhesion promoter was used. Surface control is done using a Scanning Electron Microscopy (MEB Hitachi S-4800). Finally, a perfluorodecyltrichlorosilane (FDTS) self-assembled monolayer is grafted onto the surface to prevent cells adhesion.

### Cell culture and spheroid formation

A338 cell line^15^ derived from a murine pancreatic tumor with an activating mutation of KRas oncogene (KRas^G12D^) are culture in DMEM (Sigma-Aldrich) supplemented with 10% SVF (Sigma-Aldrich) and 1% Penicillin-Streptomycin (Sigma-Aldrich), at 37°C and 5% CO_2_. Spheroid are formed using hanging droplet protocol. Typically, 15μL droplets of a cell suspension (at approximatively 13cells/μL) are dropped on a petri dish cover. To limit evaporation, 7mL of PBS are placed on the other cover part. Spheroids of 100 μm in diameter are formed in two days. In this study, we transfected PIP-FUCCI into mouse pancreatic cancer cells (A338), and used HeLa transfected with 40nm-GEMs (Genetically Encoded Multimeric nanoparticles) as in^38^.

### Agarose confinement experiments

A 48-well plate is placed on ice. We prepare a low-melting agarose solution of 2% concentration and leave it at 37°C to thermalize. 200 μL of medium containing the spheroid of 2/3 days old is then mixed with 200 μL of 2% low-melting agarose within the pipette. The 400 μL solution is placed on the 48-well plate on ice, to enable rapid polymerization of agarose at a final concentration of 1%. We find that this step is necessary to obtain a fully-embedded spheroid: if the polymerization occurs at room temperature, the spheroid sediments most of the time at the bottom of the well, and is not embedded in 3D.

### C. elegans culture

We use the *C. elegans* strain N2 (wild type), which is kindly provided by Alfonso Pérez-Escudero. *C. elegans* populations are grown, maintained, and manipulated with standard techniques^39^, except that the NGM medium is replaced by M9 agar minimal medium (M9 minimal salts supplemented with 0.2% casamino acids, 0.4% glycerol, 2.0 μg/mL thiamine and 2.5 μg/mL cholesterol). Synchronized worms are grown on agar plates seeded with a lawn of the bacteria *Ochrobactrum vermis* at 22.5 °C. Adult worms are collected in an Eppendorf tube containing 1 mL of M9 liquid medium (M9 minimal salts) and then loaded inside the microfluidic chip with a syringe. A single worm is blocked inside the chamber of the chip, grown during 48 h and fed with a unidirectional flow of a culture of *Ochrobactrum vermis* in M9 liquid, at a rate of 500 μL/h.

### D. melanogaster culture and leg preparation

Leg discs from SqhKI[RFP]3B background *D. melanogaster* are dissected at white pupal stage in Schneider’s insect medium (Sigma-Aldrich, S9895) supplemented with 15% fetal calf serum and 0.5% penicillin-streptomycin, as well as 2 μg/ml 20-hydroxyecdysone (Sigma-Aldrich, H5142). Legs are then transferred into the microfluidic chamber. Leg discs are imaged with a LSM880 confocal microscope fitted with a Fast Airyscan module (Carl Zeiss) and equipped with a 40x Water NA-1.2 objective. Stacks of 150 images with z-step of 1μm are taken every 30 minutes, with a pixel size of 0.0171μm/pixel. The laser power is set at 1%. Airyscan Z-stacks are processed through the ZEN software. Max projection images are computed and displayed on Fig. 2.

### Loading spheroids and other organisms

First, the chip is filled with DMEM medium supplemented with 10% SVF and 1% Penicillin-Streptomycin. Then, the sliding element is inserted carefully in the device to such that the cavities are aligned in front of the culture chambers. Spheroids and organisms are taken one by one using a tubing connected to a syringe. Their injection is done at the inlet localized on the side of the sliding element channel. Once a spheroid is in the channel, it will go through the sliding element and will enter the desired chamber for the high-throughput device, or the only chamber for the chemostat. This step is repeated until all the culture chambers are filled with spheroids for the high-throughput device. Then, the sliding element is moved so that each chamber is closed with a wall, or aligned with the slits / holes for feeding. The medium channel is connected to a syringe pump and a flow of 400μL/h is applied.

### Imaging conditions

A Zeiss observer microscope is used to perform the acquisition during several days. Biological samples were observed through a 63x objective. In bright-field, the exposure-time was about 100ms with 30% intensity. The environment is fixed at 37°C with 5% CO2 during the whole experiment thanks to a small incubator (tokai-hit).

### Finite element simulations

Geometry of the microfluidic cages including pressure sensor is simulated using Comsol multiphysics software with the solid mechanics module in stationary conditions. Once the geometry of the chamber is created, PDMS (polydimethylsiloxane) is set as a linear elastic material characterized by a Young’s modulus of 2 MPa, a Poisson coefficient of 0.49 and a density of 970kg/m^3^. Concerning boundaries conditions, the pressure is applied on the chamber walls which are all free to deform. Finally, a mesh controlled by the physics is applied on the structure and built with tetrahedrons elements. For each applied pressure, the total displacement of the membrane is calculated. A calibration curve describing the deformation as a function of pressure is used to calibrate all the experiments.

### Surface tension measurement

During the buildup of growth-induced pressure, the aggregate morphs from a spheroid shape to a cuboid, where the curvature decreases from the radius of the spheroid to the radius of a cell, at a given mechanical pressure. Denoting 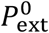 the external pressure, *P*_int_ the internal pressure, *R* the radius of curvature and *γ* the surface tension, the Laplace pressure equation can be written

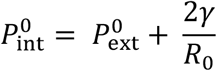

when the aggregate is a sphere, with *R*_0_ its radius, and

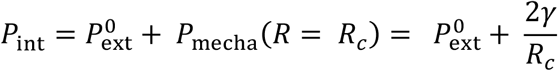

when the spheroid has morphed into a cuboid shape with curvature radius *R*_*c*_ which corresponds to the radius of a cell, and *P*_mecha_(*R* = *R*_*c*_) the mechanical pressure at this time point. *P*_mecha_(*R* = *R*_*c*_) is the pressure measured by the pressure sensor. At this surface, the curvature of the spheroid is ∼ 0 μm^-1^, the spheroid flattening on the sensor. Given that *P*_mecha_(*R* = *R*_*c*_) ∼ 300Pa, and *R*_*c*_ ∼ 10*μ*m, one gets *γ* ∼ 1.5 mN/m as a surface tension value.

### Genetically-encoded multimeric nanoparticles imaging and diffusion analysis

Experiments are performed on a Leica DM IRB microscope with spinning-disk confocal (Yokogawa CSU-X1) with a nominal power of 100mW and a Hamamatsu sCMOS camera (Orca flash 4.0 C13440) with a 63x objective. GEM nanoparticle movies are acquired by illumination with a 488 nm laser at full power. 30 images are acquired with no delay during 300 ms continual exposure at 100 Hz frame-rate. Particle tracking is achieved with the FIJI MOSAIC Suite and analyzed with a home-made Matlab script.

## Supplementary information

### Title of supplementary videos

***Video S1 –*** Growth of a multicellular spheroid in the microfluidic chemostat

[link to video](https://cloud.laas.fr/index.php/s/AI0VnnoxDCgnROk)

***Video S2 –*** Motion of the nematode *C. elegans* in the microfluidic chemostat

[link to video](https://cloud.laas.fr/index.php/s/A4hteIjk3CXZ7li)

***Video S3 –*** Development of a drosophila leg in the microfluidic chemostat

[link to video](https://cloud.laas.fr/index.php/s/r7rOkLnc5uU02o4)

***Video S4 –*** Loading of a spheroid in the confining chambers through the sliding element

[link to video](https://cloud.laas.fr/index.php/s/qDSXeSPJM8TkuLz)

***Video S5 –*** Confined growth of a spheroid and deformation of the suspended membrane with mechanical growth-induced pressure

[link to video](https://cloud.laas.fr/index.php/s/jXLISGU8a1Gddmj)

## Supplementary figures

**Figure S1:**
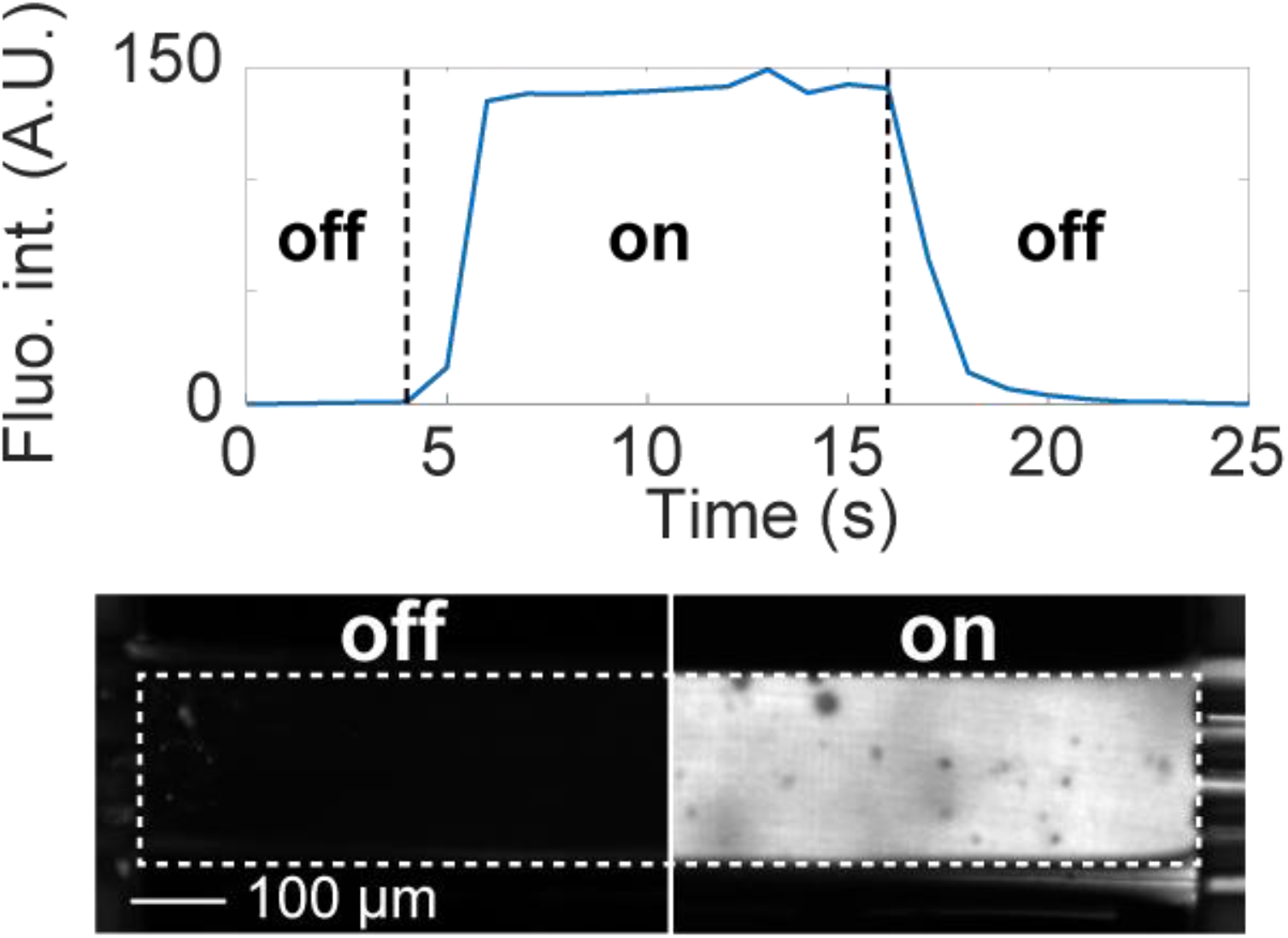
Changing of culture medium inside the device can be achieved within seconds.

**Figure S2.**
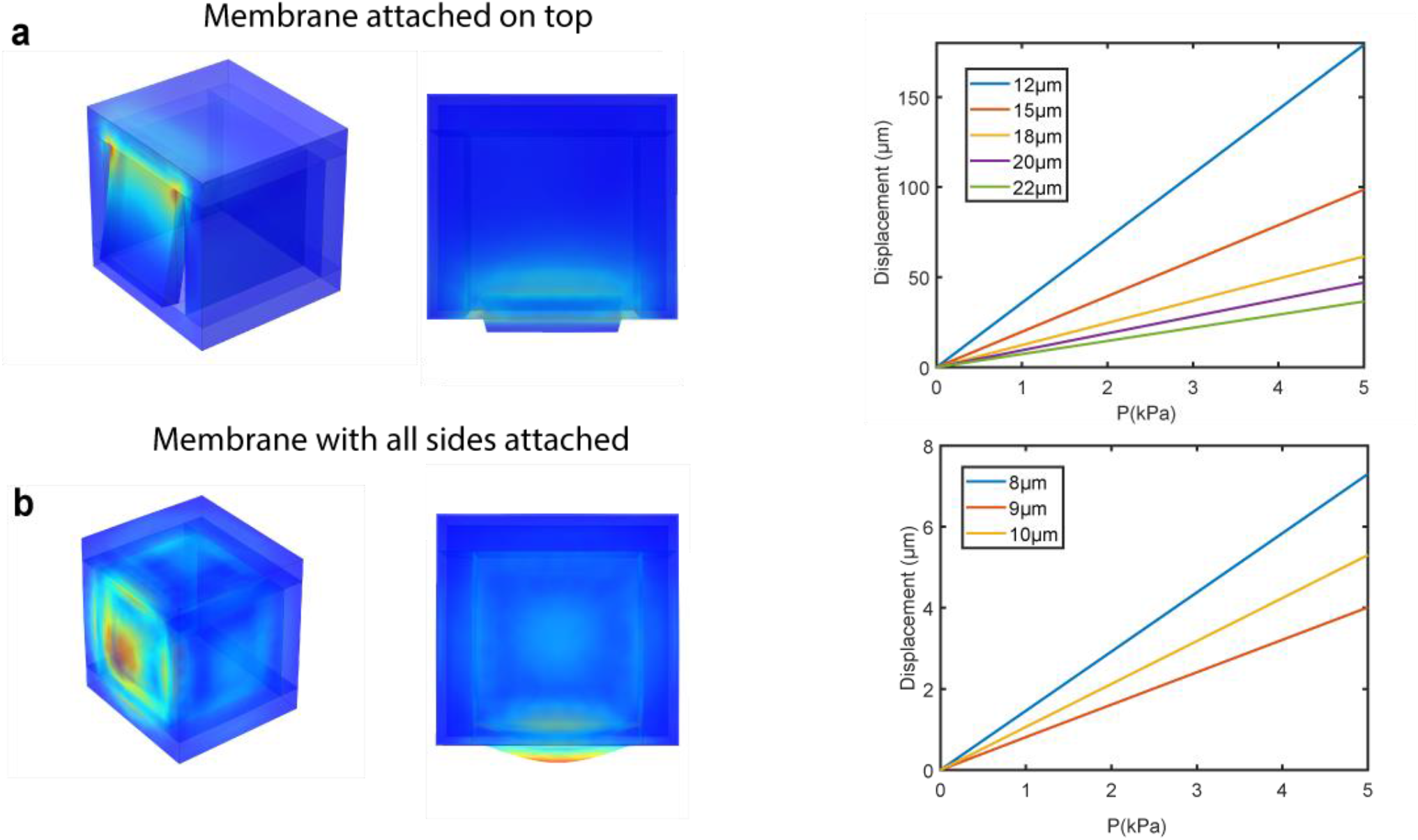
Finite element simulation of different membrane configurations to measure growth-induced pressure. **a**. Membrane only attached at the top, and **b**. membrane attached to the four sides. We notice the much higher deformability of the membrane only attached at the top.

**Figure S3:**
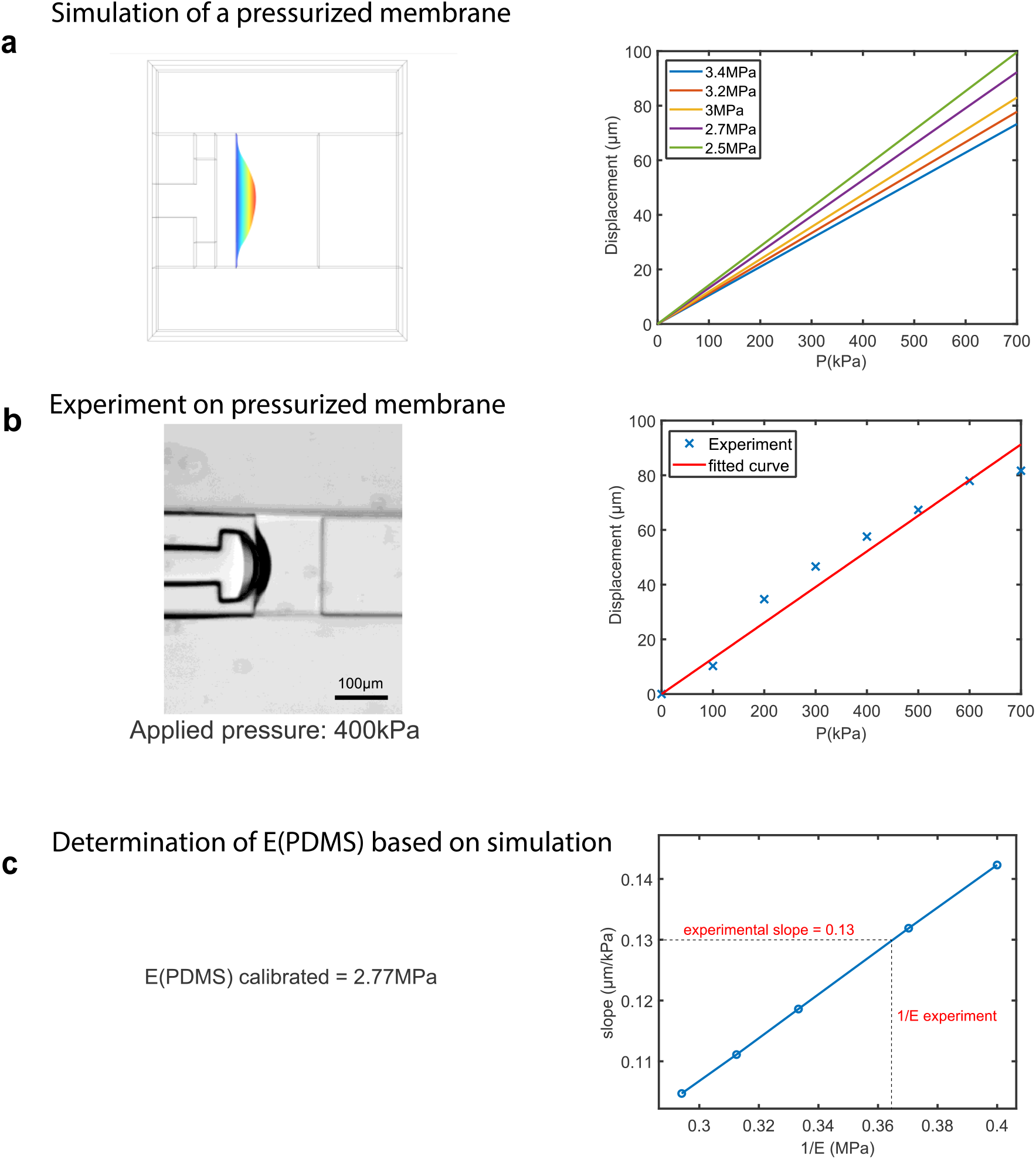
Calibration of the mechanical properties of the PDMS to use the pressure sensor. **a**. Simulation and displacement of membrane attached to its four sides as a function of the pressure for different Young’s moduli of the material. **b**. Experiment using a membrane attached to its four sides, and its deformation as a function of imposed pressure. **c**. The slope of the deformation of the simulated membrane is inversely proportional to the Young’s modulus. We use the simulation to infer the experimental Young’s modulus, and use this information together with Fig. S1 to measure growth-induced pressure.

**Figure S4:**
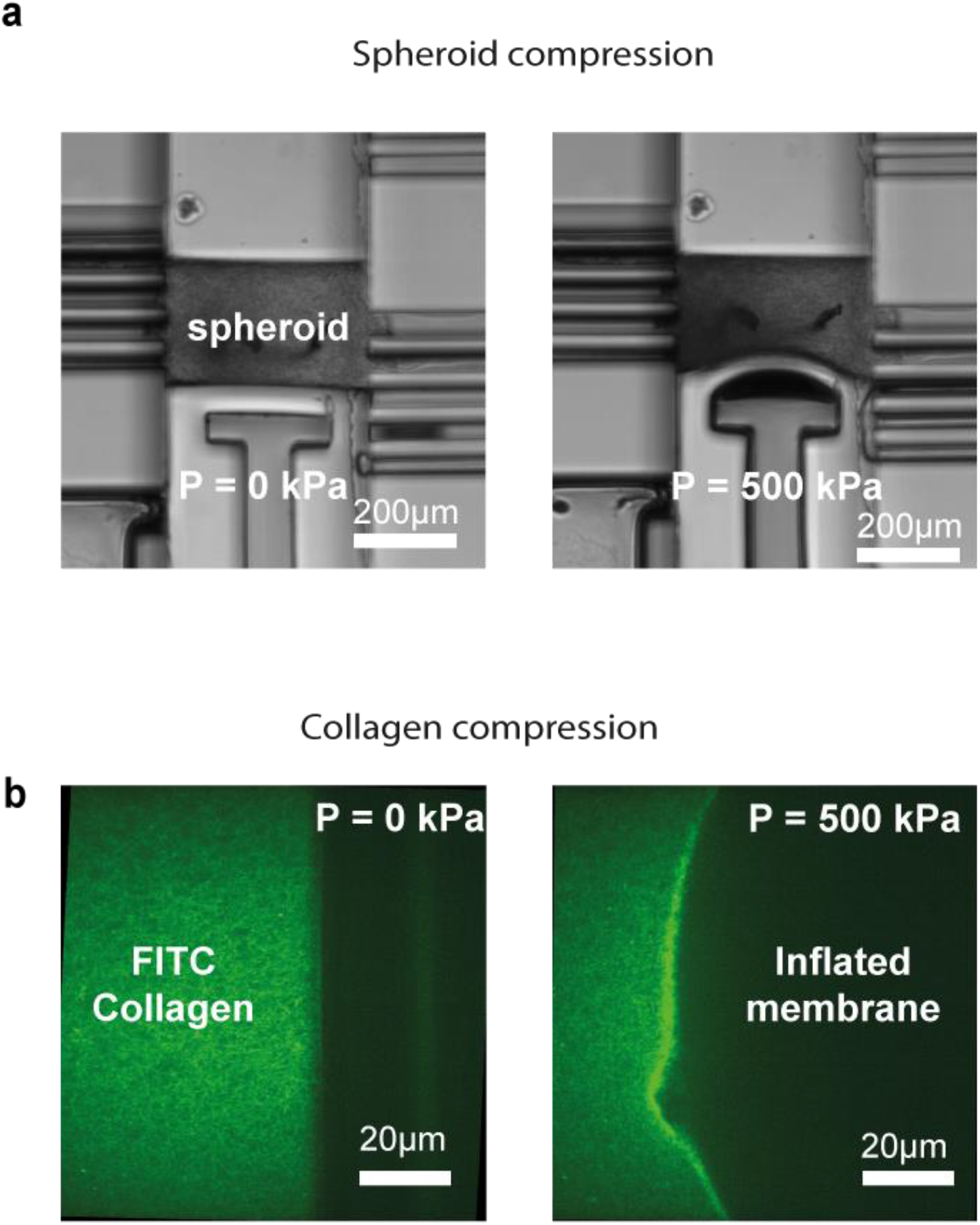
Spheroid and hydrogel compression. Using the membrane attached to every sides, we can impose a give compression onto a loaded sample, either a spheroid (**a**.) or a collagen hydrogel (**b**.).

**Figure S5:**
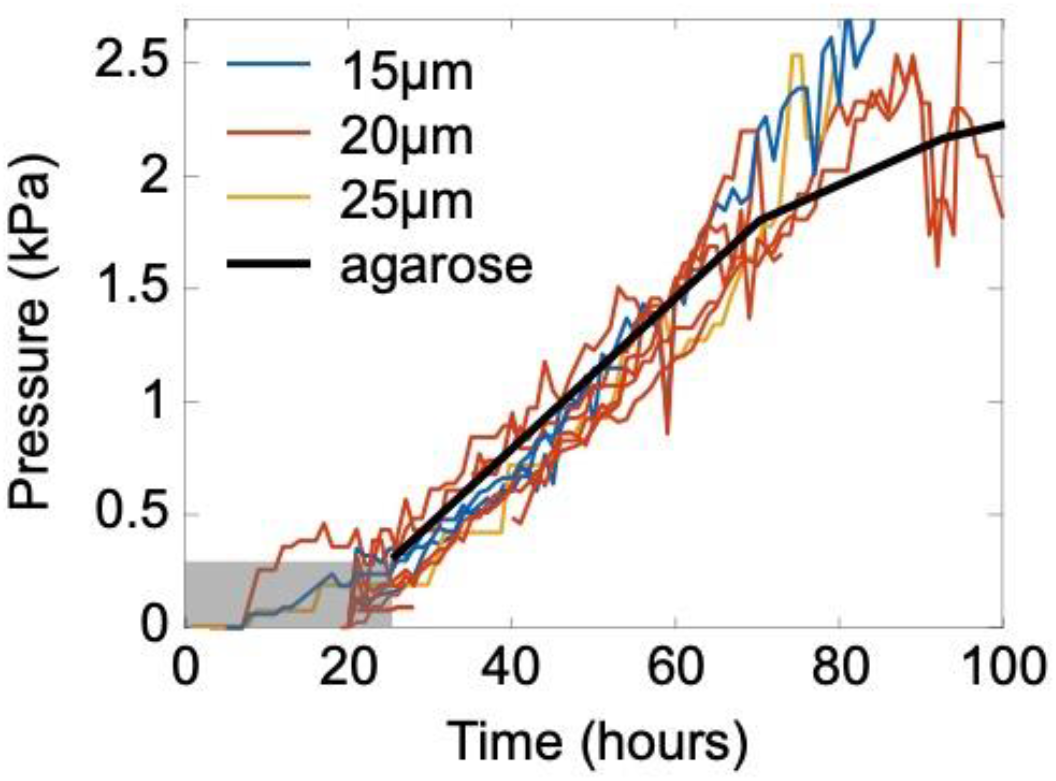
Agarose confined growth vs. microfluidic confinement. After the deformation of the spheroid to contact the whole surface of the microfluidic chamber, the spheroid is fully confined. This situation is then comparable to the case where the spheroid is fully embedded as a sphere in agarose. We thus shifted in time (24h) and in pressure (250 Pa) the agarose curve to compare the dynamics of growth-induced pressure buildup with the microfluidic confinement, and observe a similar dynamic. A potential decrease for later points inside agarose is observed, and could potentially be attributed to lesser feeding, the spheroid in agarose also being larger than in the chamber.

**Figure S6:**
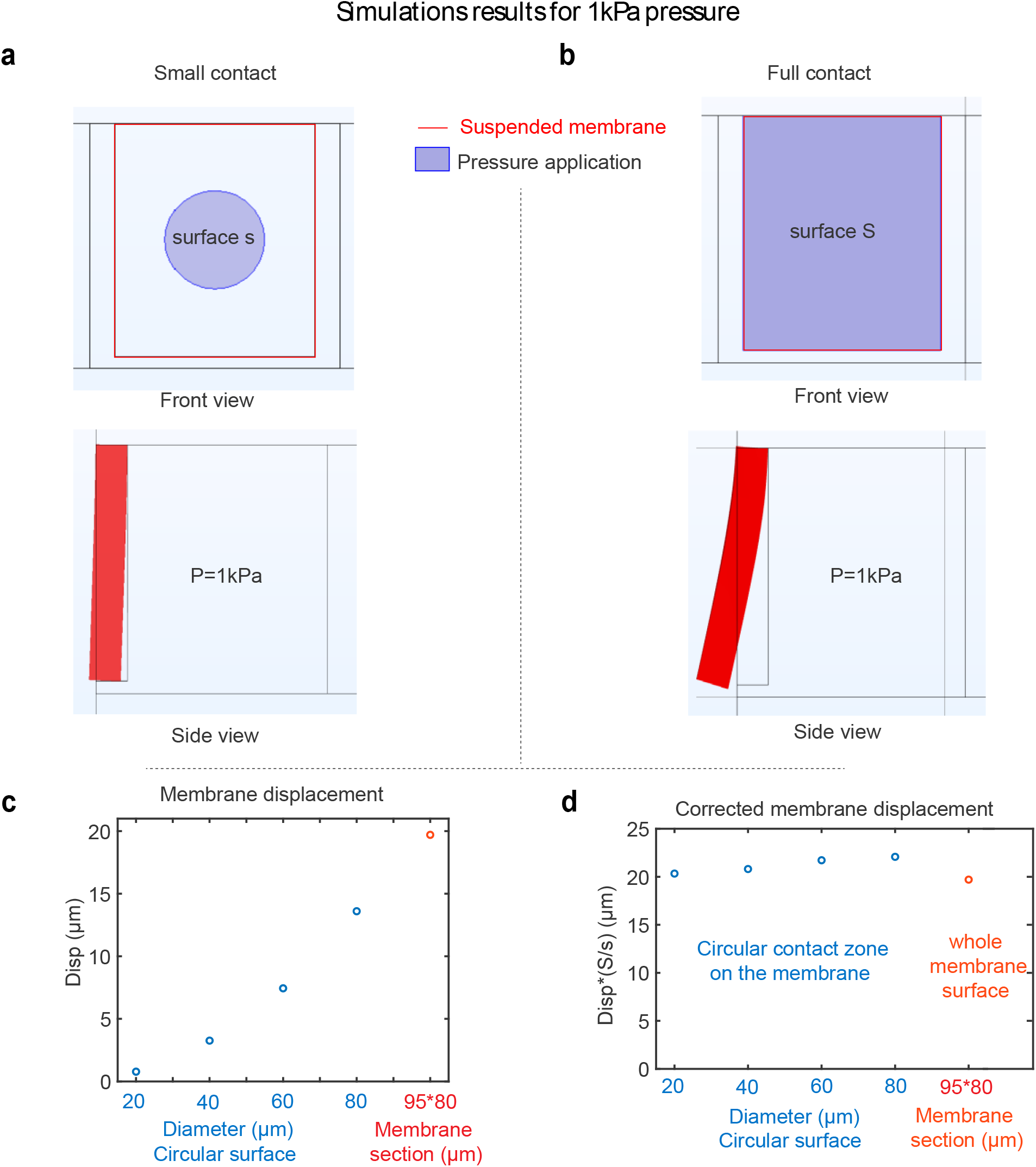
Correction factor when the spheroid does not fully contact the membrane. When the aggregate does not fully contact the surface, the pressure is applied on a smaller surface. We performed Finite Element simulations where the contact surface is either a small circle (at early time points, **a**) or fully contact the surface (at confluency, **b**). We observed that displacement increased with surface contact diameter (**c**). We showed that a correction factor of the ratio of the membrane surface to the contact surface needs to be applied (**d**). However, because the membrane does not deform uniformly, this correction factor is not exactly the ratio of the surfaces, and tends to decrease with increased contact surface.

**Figure S7:**
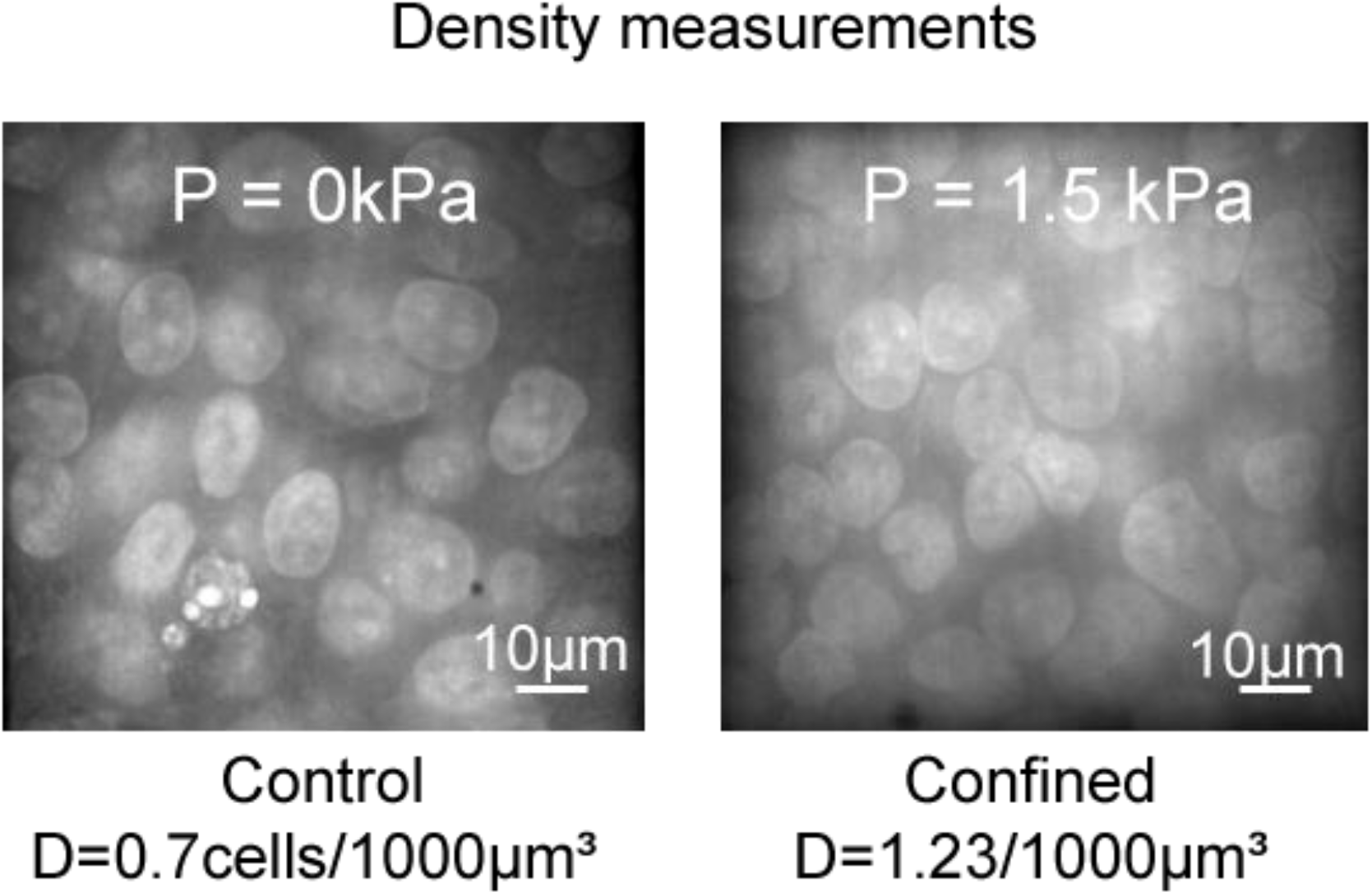
Cell density increases under confined growth. At the end of an experiment, cells were fixed and nuclei stained with DAPI. 3D stacks were taken and cell density was measured. We observe an almost doubling of cell density under an increase in growth-induced pressure.

